# Transcriptional profile of ribosome-associated quality control components and their associated phenotypes in mammalian cells

**DOI:** 10.1101/2023.02.06.527332

**Authors:** Otávio Augusto Leitão Dos Santos, Rodolfo L. Carneiro, Rodrigo D. Requião, Marcelo Ribeiro-Alves, Tatiana Domitrovic, Fernando L. Palhano

## Abstract

During protein synthesis, organisms detect translation defects that induce ribosome stalling and result in protein aggregation. The Ribosome-associated Quality Control (RQC) complex, comprising TCF25, LTN1, and NEMF, is responsible for identifying incomplete protein products from unproductive translation events, targeting them for degradation. Though RQC disruption causes adverse effects on vertebrate neurons, data regarding mRNA/protein expression and regulation across tissues are lacking. Employing high-throughput methods, we analyzed public datasets to explore RQC gene expression and phenotypes. Our findings revealed a widespread expression of RQC components in human tissues; however, silencing of RQC yielded only mild negative effects on cell growth. Notably, TCF25 exhibited elevated mRNA levels that were not reflected in protein content. We experimentally demonstrated that this disparity arises from post-translational protein degradation by the proteasome. Additionally, we observed that cellular aging marginally influences RQC expression, leading to reduced mRNA levels in specific tissues. Our results suggest the necessity of RQC expression in all mammalian tissues. Nevertheless, when RQC falters, alternative mechanisms seem to compensate, ensuring cell survival under non-stress conditions.

## Introduction

Protein synthesis is a highly regulated cellular process necessary for maintaining cellular homeostasis^1^. However, translation is prone to different errors that can cause ribosome stalling followed by ribosome collisions and premature termination. Stalled ribosomes can be a source of aberrant misfolded and dysfunctional polypeptides, which can aggregate and disrupt cellular homeostasis. In addition, stalled ribosomes are unable to initiate new translation cycles, affecting the entire dynamics of protein synthesis^2^. It was estimated that ribosome stalling occurs in 0.4% of the translation events in *Escherichia coli*^3^, therefore, cells must constantly deal with defective translation by-products. Ribosome stalling occurs when the elongation process is interrupted either by the presence of a premature stop codon, or by the absence of a termination codon, which lead to the mRNA nonsense-mediated decay and non-stop decay, respectively^4–8^. Additionally, the presence of strong structural elements in the mRNA, rare codons or polylysine coding tracts are also able to impair elongation, causing ribosome stalling and the mRNA no-go decay^4,9–15^.

Data currently available in the literature indicate that the position of the stalled ribosomes on the mRNA is important to determine how these ribosomes will be detected and rescued^16^. Ribosomes stalled mRNA collides with other elongating ribosomes forming complexes with unique structural features, such as the disomes, that are two ribosomes with their 40S subunits touching each other^17–20^. Collided disomes are recognized by the E3 ligase ZNF598 (Hel2 in yeast)^17,18^. ZNF598 recruits downstream effectors in a ubiquitylation-dependent manner^13,17,18,21–23^. One effector is the ASCC3 helicase (Slh1 in yeast), which utilizes ATP to separate the ribosomal subunits^22,24^. These mRNA degradation pathways lead to the recycling of the 40S ribosomal subunit, but leave the 60S subunit with an incomplete nascent peptide occupying the exiting tunnel and a tRNA linked to the incomplete polypeptide^4,12^.

Initially identified in yeast^25^, the Ribosome-associated Quality Control (RQC) complex acts on ribosomes stalled during elongation. It is responsible for the ubiquitination and extraction of the nascent incomplete peptide, which also leads the 60S subunit recycling. The RQC complex comprises the proteins LTN1, NEMF, and TCF25, which interact with each other and the 60S subunit bound to uncompleted polypeptides^25,26^. In the RQC pathway, ribosomes stalled and collided during translation are first sensed by factors that eventually promote subunit dissociation, a process mediated mainly by ZNF598^21^. ZNF598 is an E3 ligase that recognizes the unique structural architecture at the collision interface and ubiquitinates the 40S subunit, which leads to subunit dissociation and permits the association of the RQC with the 60S subunit^17,18,21,22,27^. After ribosome dissociation, NEMF binds to the aberrant 60S subunit contacting the polypeptide-bound tRNA, a conformation that confers specificity to aberrant 60S and prevents 40S subunit reassociation^16^. Moreover, NEMF (Rqc2 in yeast) promotes the mRNA-independent synthesis of the C-terminal Alanine Threonine (CAT) tail^28^. This CAT tail has two distinct functions: it is a stress signal due to its hydrophobic nature, and it assist the ubiquitination of the nascent peptide through the exposure of lysine residues that may be enclosed inside the ribosomal exit tunnel^25,29^. Following NEMF biding to the 60S, the ubiquitination of the nascent peptide is performed by the E3 ligase LTN1, a process that apparently depends on TCF25^30,31^. The molecular function of TCF25 is still not fully understood, but as previously shown by our group, the expression of its yeast homolog, Rqc1, is regulated post-translationally by LTN1, perhaps as a negative feedback mechanism to ensure that the RQC complex is not over activated^15^. It is worth noting that TCF25 was originally described as a transcriptional factor, and it was recently implicated in the expression regulation of NFAT (Nuclear factor of activated T-cells) in cardiomyocytes and the activation of T-cells during papillary renal cell carcinoma^32–35^.

Defects in the RQC complex have been associated with neurodegeneration and protein aggregation. Chu et al., 2009, demonstrated that homozygous LTN1 knockout mice exhibit age-dependent loss of locomotor ability. Besides, pathological analyses evidenced dystrophic neurites, gliosis, vacuolated mitochondria, and accumulation of soluble hyperphosphorylated tau in these mice. Although LTN1 is widely expressed in all tissues, motor and sensory neurons and neuronal processes were the most affected^36^. Choe et al., 2016, showed that deletion of the LTN1 in yeast causes stalled nascent peptides to form detergent-resistant aggregates and inclusions. This aggregation process depended on CAT tail formation by Rqc2, while the double knockout ΔLTN1ΔRqc2 prevented the formation of these aggregates in yeast^37^.

Similarly, CAT tailing by NEMF in HeLa cells can form aggregates when LTN1 is depleted^38^. The dysfunction of NEMF is also associated with neurological disorders. Martin et al., 2020, reported that mice with NEMF mutations R86S and R487G present many neuromuscular phenotypes. Moreover, R86S mice died prematurely, with a median lifespan of 20 days. Furthermore, it was already observed that people with mutations in NEMF display intellectual disability, speech delay, neuropathy, and motor dysfunction with varying severities and progression^39^.

Recently, it was shown that aged *Saccharomyces cerevisiae* and *Caenorhabditis elegans* under chronological aging downregulate the mRNA of some RQC components and have increased ribosome stalling and collisions at polybasic tracts^40^. It was proposed that the RQC activity is saturated by an age-dependent increment in ribosome stalling events, which could lead to impaired proteostasis observed in older individuals. However, it was not demonstrated that aging affects the activity and expression of RQC components in mammalian tissues.

The above evidence indicates that neuronal cells physiology is particularly affected when the RQC complex is perturbed. It is unclear, however, whether the expression of the components of the RQC varies across different tissues. Also, it is not known if the mRNA levels of the RQC components are co-regulated, and if aging perturbs the expression. We took advantage of publicly available results from high-throughput experiments such as transcriptomes, proteomes, pertubseq and others listed in **Table 1** to map various aspects of RQC components expression and regulation. Even though ZNF598 is not regarded as an integrant part of the RQC complex^18^, for the sake of simplicity, it will be referred as an RQC component from now on.

**Table 1:**
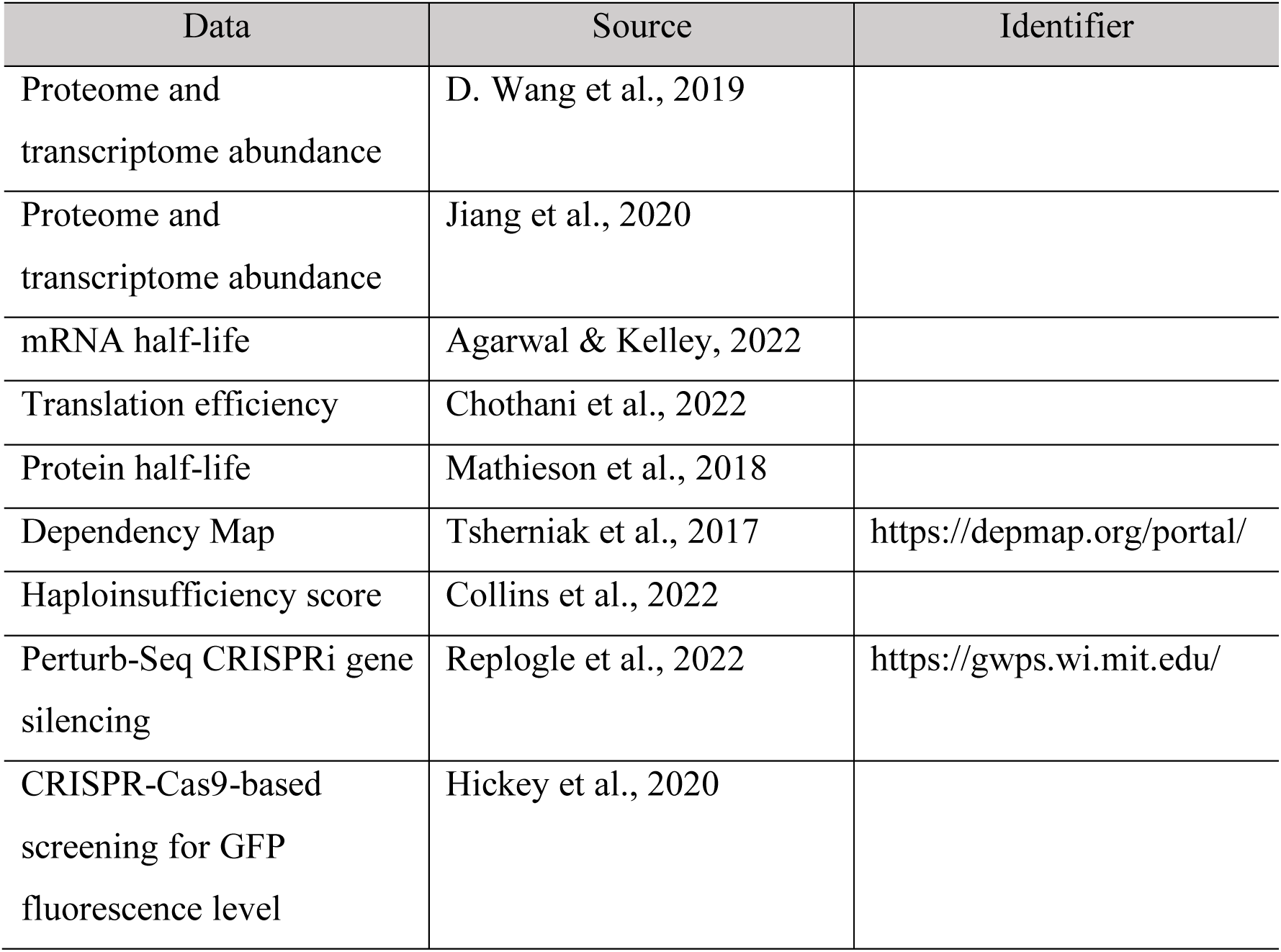
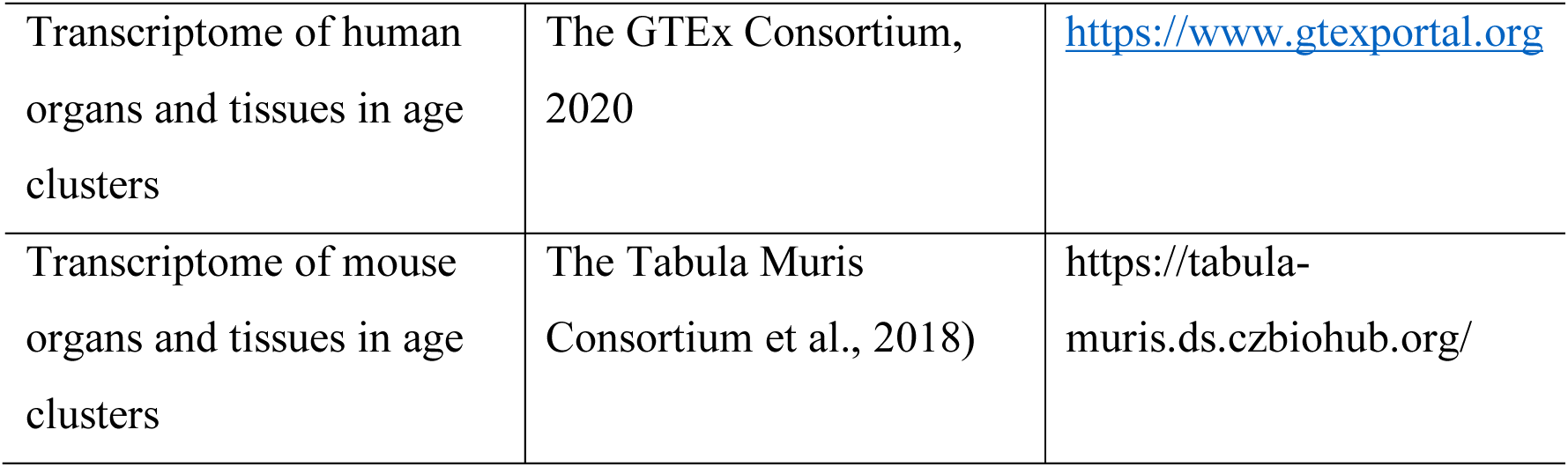
Public data assessed in the study.

| Data | Source | Identifier |
| --- | --- | --- |
| Proteome and transcriptome abundance | D. Wang et al., 2019 |  |
| Proteome and transcriptome abundance | Jiang et al., 2020 |  |
| mRNA half-life | Agarwal & Kelley, 2022 |  |
| Translation efficiency | Chothani et al., 2022 |  |
| Protein half-life | Mathieson et al., 2018 |  |
| Dependency Map | Tsherniak et al., 2017 | https://depmap.org/portal/ |
| Haploinsufficiency score | Collins et al., 2022 |  |
| Perturb-Seq CRISPRi gene silencing | Replogle et al., 2022 | https://gwps.wi.mit.edu/ |
| CRISPR-Cas9-based screening for GFP fluorescence level | Hickey et al., 2020 |  |
| Transcriptome of human organs and tissues in age clusters | The GTEx Consortium, 2020 | https://www.gtexportal.org |
| Transcriptome of mouse organs and tissues in age clusters | The Tabula Muris Consortium et al., 2018) | https://tabula-muris.ds.czbiohub.org/ |

We found that all RQC components are ubiquitously expressed in the analyzed tissues. Interestingly, TCF25 showed a high number of mRNA transcripts, which does not translate into high protein levels in most tissues analyzed. In addition, when compared to the rest of the genome, TCF25 had increased mRNA half-lives coupled with shorter protein half-lives, suggesting a post-translational regulation. To experimentally test this observation, we checked TCF25 expression in HEK cells by western-blot and observed that TCF25 protein levels can be increased by inhibiting the proteasome, what indicates that a post-translational regulation determines the TCF25 expression level. By analyzing published pertub-seq data, we observed that silencing RQC components caused only mild effects at normal growth conditions in different cell lines. Finally, we dissected the effect of aging on the mRNA levels of the RQC components in humans and mice. In general, aging caused a discrete decrease in mRNA levels in some human tissues. Moreover, we found that LTN1 and NEMF expression levels and associated phenotypes tended to be similar, while TCF25 and ZNF598 had specific, uncorrelated patterns. In conclusion, our results point out that the RQC components are ubiquitously expressed in mammalian tissues, and we found support for aging-dependent downregulation that might regulate RQC function. Additionally, although TCF25 interacts with LTN1 and NEMF, it is subjected to different expression regulation.

## Materials and Methods

### Transcriptome and translatome profile

To assess the transcription and translation profile of the components of the RQC complex, we obtained data from D. Wang et al., 2019 obtained from 29 healthy human tissue samples. For each RQC complex component (LTN1, TCF25, NEMF, and ZNF598), the transcripts per million (TPM) values were used to estimate the Z-score for each gene across the tissue and we plotted the result as a heatmap. To study the proteome in each tissue, we used the dataset containing relative protein abundances (log_2_) and estimated the Z-score for each gene. A similar approach was performed with Jiang et al., 2020 dataset in the GTEx Consortium. From this data, we obtained the transcriptome and proteome of 114 samples from 32 organs from 14 individuals, with replicates ranging from 1 to 22.

### RNA and protein half-life

To assess the half-life of transcripts for each gene in the RQC complex, we used the dataset produced by Agarwal & Kelley, 2022. They generated a compendium of mammalian (mouse and human) mRNA half-life datasets from 33 publications obtained from cell culture and normalized the values using the Z-score. For protein half-life, we assessed the data from five different non-dividing cell types (human B-cells, monocytes, NK cells, hepatocytes, and mouse embryonic neurons) by Mathieson et al., 2018. We calculated mean protein half-live values considering all cells and compared them with the values for each component of the RQC complex in each cell line analyzed.

### Translational efficiency

To analyze the mRNA translation efficiency (TE), we downloaded the dataset from ribosome profiling experiments published by Chothani et al., 2022.

### Impact on growth analyses

The impact of the deletion of RQC complex components genes was obtained using the large-scale gene essentiality data from the DepMap project^46^. They performed genome-scale library of 100,000 short hairpin RNAs (shRNAs) to analyze a genome-scale loss-of-function screens performed in 1,086 human cell lines. A scoring system was created to quantify the effect of gene inactivation on cellular growth in a specific cell lineage. If the gene inactivation caused no effect in a specific cell lineage, a score equal to zero was attributed to this pair (cell line/gene inactivated). On the other side, a score of -1 meant that this pair led to cell death, and it corresponded to the median of all common essential genes. Data were obtained directly from the DepMap portal (https://depmap.org). We searched in “perturbation effects” and downloaded the dataset (DepMap 22Q2 Public+Score, Chronos) for each RQC complex gene. Furthermore, we plotted the correlation between growth score values obtained for NEMF and LTN1 silencing on different cell types.

### Haploinsufficiency score

The haploinsufficiency score was created through the meta-analysis of approximately one million individuals in both disease and control cohorts to identify disease-associated dosage-sensitive regions, allowing for dosage sensitivity for each gene and creating a haploinsufficiency score^47^. We used the score obtained for each gene of interest for comparison purposes. Genes with values equal to or greater than 0.86 were considered haploinsufficient.

### Perturb-Seq CRISPRi gene silencing

To evaluate the effects of RQC impairment on a transcriptional level, we analyzed data from Replogle et al., 2022, which performed a genome-scale disturb-seq in leukemia K562 cell line and investigated the mRNA expression profile of individually silenced genes. The data was obtained from the online platform https://gwps.wi.mit.edu/, which separates the top 30 up and downregulated genes for each selected gene. We used the K562 Genome-Wide Perturb-Seq database, selected the top 30 up and downregulated component genes of the RQC complex, and separated them into clusters to understand the transcriptional profile. Also, we used the 10 genes with the most similar transcriptional profiles provided by the tool for each RQC complex component.

### Gene ontology

We selected gene clusters (up and downregulated) that present similar transcriptional profiles between NEMF and LTN1 and performed a gene ontology analysis. The geneset enrichment ontology analyses were performed in the Gene Ontology Consortium (http://geneontology.org/) with the GO Enrichment Analysis tool, searching for “Biological process”. We selected the results with the highest Fold enrichment.

### FACS-based CRISPRi screen

To screen if the genes participate in the RQC complex, we used data from Hickey et al., 2020, which measures the expression of a fluorescence reporter lacking a stop codon (leading to non-stop decay) in a genome-wide CRISPRi screening. Positive values indicate that the knockdown of that gene stabilizes the GFP signal. We used the RQC complex component as a control.

### Age-dependent transcriptome

To assess if the transcriptional profile of RQC components changes with age, we used data from the GTEx consortium for human tissues (The GTEx Consortium, 2020) and The Tabula Muris Consortium for mouse tissues (The Tabula Muris Consortium et al., 2018). We downloaded the data from the GTEX consortium data on the website https://www.gtexportal.org for each human tissue and separated them into age clusters (20-29, 30-39, 40-49, 50-59, 60-69, and 70-79). We used the transcript per million (TPM) to determine the mRNA abundance across the human tissues at different ages clusters. The data was normalized by the reference age range (20-29). For mouse data, we used The Tabula Muris Consortium, which offers transcriptomes of 20 organs from four male and three female mice from 1 to 27 weeks old. Data handling and data presentation were similar to that of human samples.

### Proteasome and autophagy inhibition

For the proteasome inhibition and autophagy assay, we used 1 x 10^6^ cells/wells HEK 293 cells cultured in a 6-well plate. After 24 hours, the media was removed, and cells were gently washed with 1X PBS pH 7.4 and incubated with 10 µM MG-132 (proteasome inhibitor) and 50 µM chloroquine (autophagy inhibitor) at 37°C for 3 hours in the presence of 100 µg/ml cycloheximide. After this period, the culture medium was removed, 100 µl of ice-cold RIPA (Thermo Fisher) lysis buffer with PMSF 1 mM and Halt™ Protease and Phosphatase Inhibitor Cocktail 1X (Thermo Fisher) was added, and the cells were scraped. All these procedures were performed on ice. Cells were added into microtubes, lysed with a microtube macerator, and centrifuged at 9,000 rpm for 15 minutes at 4°C. Then, the supernatant was collected, added 4X sample buffer, and heated at 100 °C for 10 minutes to perform Western blotting.

### Western blotting

Proteins separated by 12.5% SDS-PAGE were transferred to PVDF membranes using BIO-RAD’s TRANS-BLOT semi-dry system in transfer buffer (25 mM Tris, 192 mM glycine, 0.1% SDS, 20% methanol, pH 8.3). Membranes were blocked using a Li-Cor blocking buffer for 2 hours at 4°C. Incubation with primary antibodies was performed at 4 °C overnight. Membranes were incubated in Li-Cor Odyssey secondary antibodies for at least 3 hours and then observed in an Odyssey scanner. The primary antibodies used were anti-Nulp1 (TCF25) mouse (Invitrogen; 1:3,000 dilution) and anti-α-actin mouse (Thermo Scientific; 1:5,000 dilution). The secondary antibody was the anti-mouse 800 CW Li-Cor (1:5,000 dilution).

## Results and Discussion

### RQC mRNA and protein expression across human tissues

To describe and compare the expression levels of the RQC components in different human tissues, we curated previously published work that generated paired transcriptome and proteome datasets of human samples. D. Wang et al., 2019, investigated the transcriptome and proteome from 29 different healthy human tissues from the Human Protein Atlas project. Using this dataset, we investigated the mRNA and protein levels of LTN1, TCF25, NEMF, and ZNF598 (**Figure 1**). Transcripts were compared by determining the Z-score for each gene using the transcripts per million (TPM) values. Protein levels derived from proteome data were compared by determining the relative protein abundance (log_2_) (**Figure 1**). All RQC components were expressed in most tissues analyzed at roughly similar levels. So, even though RQC malfunction particularly impacts the nervous system in vertebrates^36,39^, this specificity could not be explained by the expression levels of RQC components. LTN1 and ZNF598 showed lower mRNA levels, while NEMF showed a Z-score close to zero for each tissue. Surprisingly, TCF25 showed approximately twice as much mRNA levels when compared to the average gene expression in most tissues. The protein and mRNA levels showed a similar pattern for LTN1, NEMF, and ZNF598. The exception was TCF25. As mentioned above, the mRNA levels for TCF25 were higher than the average gene expression in every tissue analyzed, and it was the most abundant RQC component transcript (**Figure 1**). However, this high amount of mRNA was not reflected in protein levels, which was generally similar to the other RQC components.

**Figure 1.**
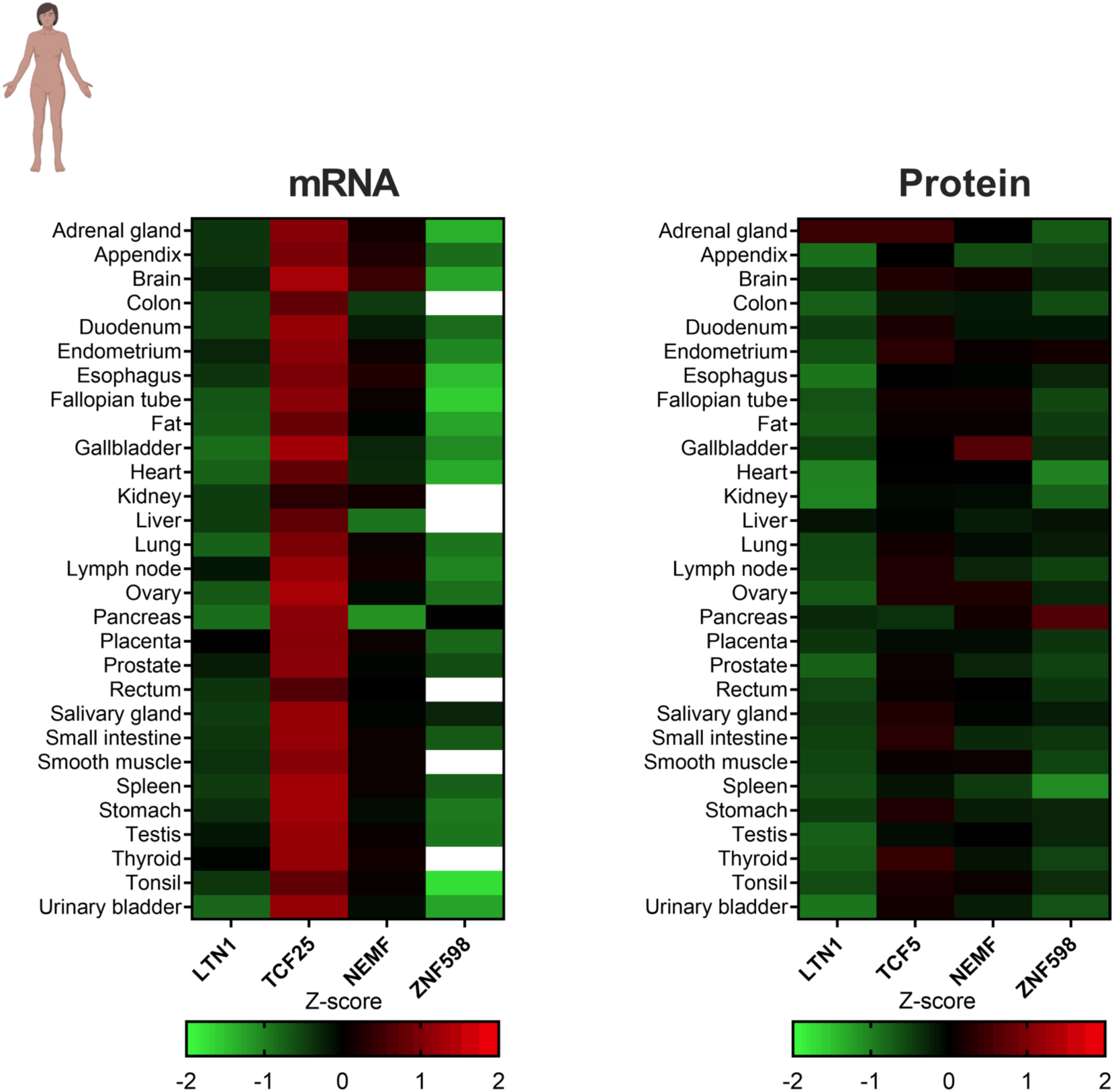
Heat map of the Z-score distribution of the RQC components calculated from the transcriptome and proteome of human tissue samples. Data obtained from D. Wang et al., 2019. White rectangles denote the absence of data.

In this study, D. Wang et al., 2019 used only one sample per tissue to generate the protein and mRNA expression data. Thus, the small sample size undermines a solid conclusion about subtle expression variations between tissues. However, despite this limitation, this dataset allowed us to observe that: 1) both mRNA and proteins of the RQC components are expressed in most tissues, 2) TCF25 mRNA is expressed in high levels, while its protein has expression levels closer to the average in most tissues (**Figure 1**).

We repeated the same analysis using another data set with a bigger sample size. Jiang et al., 2020 analyzed the transcriptome and proteome of 114 samples from 32 major organs from 14 different individuals, with biological replicates ranging from 1-22. As in Figure 1, we used the Z-score to compare the mRNA and protein levels of the RQC components (**Figure 2**). We observed that, overall, the expression levels were similar to or higher than the average of the genome (**Figure 2**). For NEMF and ZNF598, protein levels were slightly higher than the global average, showing a positive correlation with their mRNA levels. TCF25 mRNA levels were above the 75th percentile (3^rd^ quartile) in all 32 tissues studied. On the other hand, TCF25 protein expression levels were above the 75th percentile just in 10 out of 32 tissues, namely, the adrenal gland, brain cortex, minor salivary, ovary, pancreas, pituitary, prostate, thyroid, uterus, and vagina. For the other 22 tissues, the protein Z-scores were relatively lower than the mRNA Z-score (**Figure 2**).

**Figure 2.**
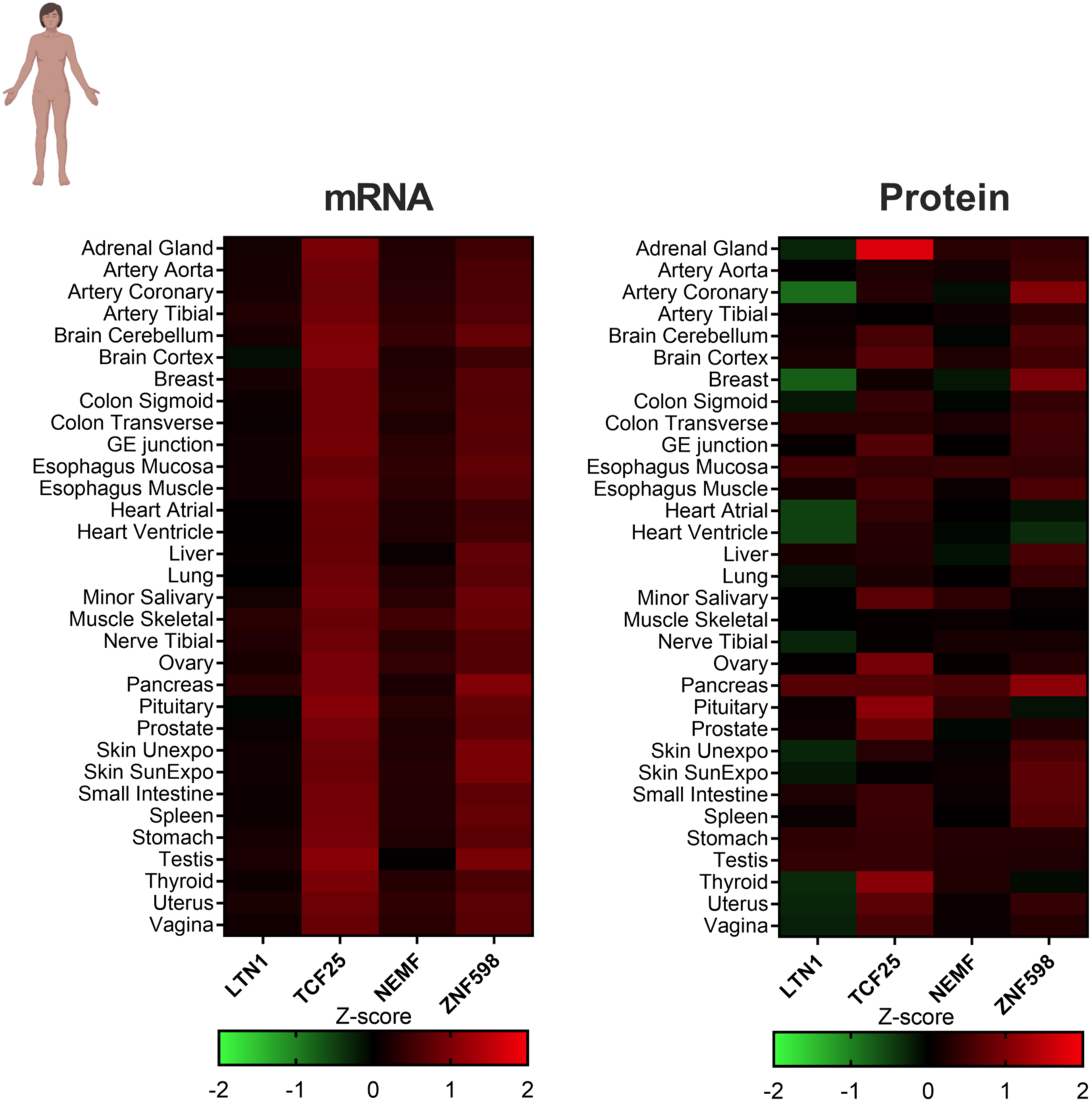
Heat map of the Z-score distribution of the RQC components calculated from the transcriptome and proteome of 114 samples of 32 major organs from 14 different individuals, with replicates ranging from 1-22. Data obtained from Jiang et al., 2020.

However, the two datasets pointed out that TCF25 protein levels are relatively low for the mRNA level detected in many tissues compared to other RQC components (**Figures 1 and 2**). This observation suggests that TCF25 may be regulated at the level of translation and/or post-translation.

The RNA transcripts are determinants for protein abundance, and, in most cases, higher mRNA levels lead to higher protein abundance^53^. However, many factors can affect the mRNA/protein ratio in a dynamic and complex process^54^. The amount of protein generated by mRNA results from different, non-mutually exclusive factors or regulatory processes such as codon usage, mRNA half-live, and protein turnover^53,55^.

To assess if TCF25’s relatively low protein levels observed in Figures 1 and 2 could be related to processes involving mRNA stability, we evaluated the half-live of the mRNA of each RQC component (**Figure 3A**). We used data produced by Agarwal & Kelley, 2022, which generated a compendium of mammalian (mouse and human) mRNA half-life datasets from 33 publications obtained from cell culture. For human cells, the mRNA half-life for LTN1, NEMF, and ZNF598 tended to be close to most transcripts in these cells (**Figure 3A dotted line**). In contrast, TCF25 showed a slightly higher mRNA half-life with mean Z-score values around 0.9. These findings indicate that the decrease in TCF25 protein levels is not due to a short half-life of its mRNA. It is important to note that while mRNA and protein quantification presented herein were obtained from studies using human tissues (**Figures 1 and 2**), the mRNA half-life data were obtained using cultured cells (**Figure 3A**).

**Figure 3.**
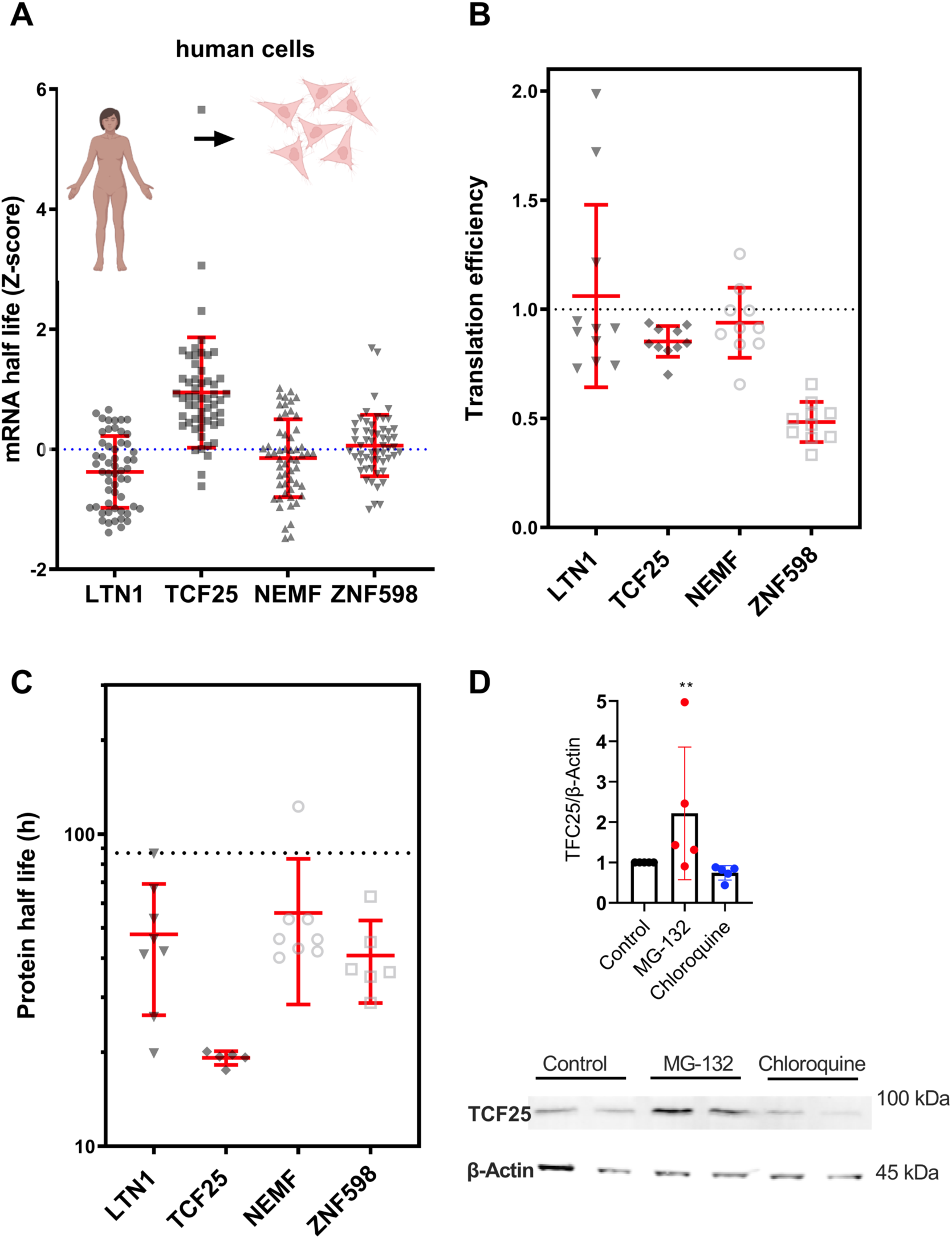
Factors that affect the abundance of RQC complex components. (**A**) Z-score calculated for the half-life of RQC components transcripts in human cells. Data obtained from Agarwal & Kelley, 2022. (**B**) Log_2_ of the mRNA translation efficiency of each transcript from six primary human cell types and five tissues. Data obtained from Chothani et al., 2022. (**C**) Protein half-life for each RQC component. The dotted line refers to the average half-life for all proteins considering all cell lines tested (86.95 hours).

Another factor that can modulate the protein amount produced by mRNA is the translation efficiency (TE). To investigate the effect of TE on TCF25 expression, we used a dataset from six primary human cell types and five tissues published by Chothani et al., 2022. They determined the translation elongation rates for the entire transcriptome through ribosome profiling. The TE values showed that TCF25 accompanied NEMF and LTN1 while ZNF598 presents lower TE values (**Figure 3B)**. Therefore, the translation efficiency is probably not the factor behind the relatively low TCF25 protein levels. We investigated the protein half-life and turnover using data from five different non-dividing cell types (human B-cells, monocytes, NK cells, hepatocytes, and mouse embryonic neurons) by Mathieson et al., 2018. All RQC components and ZNF598 showed a shorter half-life than the average, but TCF25 showed the lowest half-life with less than 20 hours for all cell types tested (**Figure 3C)**. As the transcriptome and proteome analysis suggested that TCF25 protein has a fast turnover, we decided to check if TCF25 levels are controlled by any of the two main pathways for protein degradation: the proteasome and autophagic vesicles. For this, HEK293 cells were treated with cycloheximide to block protein synthesis, and MG132 or chloroquine was added to inhibit the proteasome or autophagy, respectively. After a 3 hours treatment **(Figure 3D)**, we measured the TCF25 levels by western blot. We detected an increase in TCF25 after proteasome inhibition when compared with the vehicle control. No difference was observed when autophagy was inhibited **(Figure 3D)**. We concluded that, at least for HEK293 cells, the TCF25 protein levels are regulated by the proteasome.

Data obtained from Mathieson et al., 2018. (**D**) Western blot of TCF25 from HEK293 cells treated for 3 hours with 100 µg/ml cycloheximide to block protein synthesis. 10 µM MG132 or 50 µM chloroquine was added to block the proteasome or autophagy. DMSO was the control vehicle and β-actin was measured as loading control. ** p = 0.0085 by Friedman’s test.

### The effect of RQC components disruption in tissue cells survival and gene expression profiling

We saw that the components of the RQC complex are ubiquitously expressed in mammals, which indicates their housekeeping role. Besides the well-established co-translational quality control function described above, RQC components can be implicated in other processes, as in the case of TCF25. Nevertheless, gene deletion of RQC components showed mixed results depending on the organism. Deletion of LTN1 has no detectable effect in yeast cells growing in optimal condition, and double knockout of LTN1 and Rqc1 are also viable^15,25^. These observations suggest that activation of the RQC complex may be only necessary for specific stress situations. On the other hand, knockout mice for LTN1 exhibit strong neuromotor phenotype, and NEMF mutations in humans and mice also lead to neuromotor impairment^36,39^, which points that the RQC is necessary during neuronal development.

To further examine to what extent the RQC genes are fundamental components of living cells, we utilized the large-scale gene essentiality data from the DepMap project^46^. A scoring system was created to quantify the effect of gene inactivation on cellular growth in a specific cell lineage^46^. If the gene inactivation caused no effect on cellular growth in a specific cell lineage, a score equal to zero **(dotted black line, Figure 4A)** was attributed to this pair (cell line/gene inactivated). On the other side, a score of -1 **(blue dotted line, Figure 4A)** meant that this pair led to cell death and corresponds to the median of all common essential genes. The lowest score for the RQC components was -0.7, meaning that they are not essential for the *in vitro* growth of mammalian cells (**Figure 4A**). However, the median score for all RQC components analyzed was below zero, suggesting that the perturbation of RQC causes some negative effect on cellular growth (**Figure 4A**).

**Figure 4.**
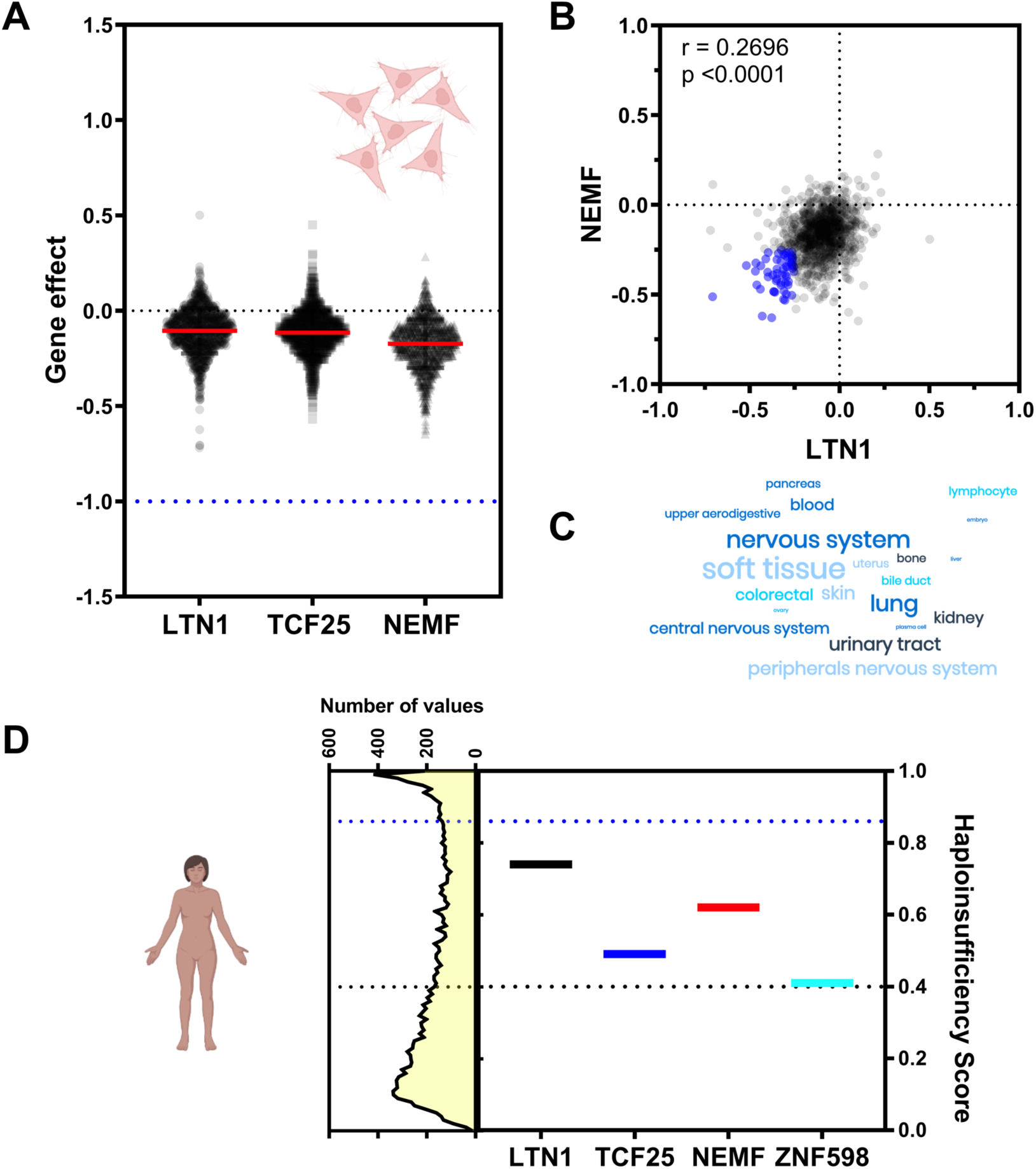
The genes of the components of the RQC complex are not essential. (**A**) Score attributed to the impact of gene inactivation on growth rate in 1086 cancer cell lines for LTN1, NEMF, and TCF25. Data obtained from Tsherniak et al., 2017. (**B**) Correlation graph of silencing effect between NEMF and LTN1. (**C**) Word cloud representation of the cellular lineages is more affected by NEMF and LTN1 silencing. (**D**) Haploinsufficiency score for each component of the RQC complex and ZNF598. Genes with values equal to or greater than 0.86 are considered haploinsufficient. Data obtained from Collins et al., 2022.

Another feature extracted from this dataset is the correlation between growth score values obtained with different silenced genes on different cell types. We found a positive correlation between NEMF and LTN1, meaning that the silencing of these genes causes a similar effect on growth depending on the cellular lineage analyzed (**Figure 4B**). To see which cellular lineages are more affected by NEMF and LTN1 silencing, we arbitrarily defined a cutoff of -0.25 (**blue circles of Figure 4B**) for both genes. Soft tissue and nervous system cellular lineages were the most affected by NEMF and LTN1 silencing (**Figure 4C**), which agrees with the neurological impairments observed *in vivo*^36,39^.

Another important consideration is to assess the haploinsufficiency of the RQC genes. Haploinsufficiency is defined as a dominant gene phenotype in diploid organisms, in which a single copy of the wild-type allele at a locus in a heterozygous combination with a variant allele is insufficient to produce the wild-type phenotype. In this context, we evaluated whether defects in one of the alleles of the genes of the RQC complex would be sufficient to alter the wild-type phenotype and thus correlate with the growth impairment observed in the knockout cells **(Figure 4C)**. In addition, recent work performed a meta-analysis of approximately one million human individuals in disease and control cohorts to identify disease-associated dosage-sensitive regions^47^. The authors predicted dosage sensitivity for each gene, creating a haploinsufficiency score. In other words, a score value equal to or higher than 0.86 (**blue dotted line, Figure 4D**) meant that a single copy of the wild-type allele at a locus in a heterozygous combination with a variant allele was insufficient to produce the wild-type phenotype. When we analyzed the haploinsufficiency score of the RQC components, we observed that all genes had a score lower than 0.86, meaning that these genes are not haploinsufficient because just a single copy of the wild-type allele is enough to produce the wild-type phenotype (**Figure 4D**). This data agrees with the description of families with pathogenic biallelic NEMF variants with a spectrum of central neurological disorders^39^. In conclusion, in vertebrate cellular lineages, knockdown of the RQC components causes a mild phenotype that can be exacerbated in specific tissues (**Figure 4A**). This result suggests that even though the RQC components are ubiquitously expressed genes (**Figures 1 and 2**), the cell can tolerate the translational perturbations caused by RQC impairment and keep growing, perhaps by adapting the transcriptional program. It is important to note that this scenario might be different under stress conditions.

### The effect of RQC components disruption in tissue cells transcriptome

To understand how cells respond to RQC impairment on a transcriptional level, we analyzed data from Replogle et al., 2022, which performed a genome-scale CRISPRi disturb-seq in leukemia K562 cell line to determine the mRNA expression profile of individually knockdown genes. The transcriptional responses were compared with the control cell. Using these data, we separated the top 30 up and downregulated genes of the RQC components in clusters **(Figure 5A)**. As expected, LTN1 and NEMF showed similarities in both up and downregulated genes, but the silencing of TCF25 showed different patterns (**Figure 5A**). These results further support that TCF25 has additional functions other than co-translational control, as the transcriptional regulation described above, and additional studies are necessary to evaluate which genes are being regulated directly by this protein. We performed an ontology enrichment analysis of the gene clusters that showed the greatest similarity in expression profile when we silenced LTN1 and NEMF. The enriched ontologies of the upregulated gene clusters were involved in negative regulation of the ubiquitination process and ribosome assembly (**Figure 5B**). In contrast, the enriched ontologies for the downregulated gene clusters were involved in the electron transport chain and free ubiquitin chain polymerization (**Figure 5B**).

**Figure 5.**
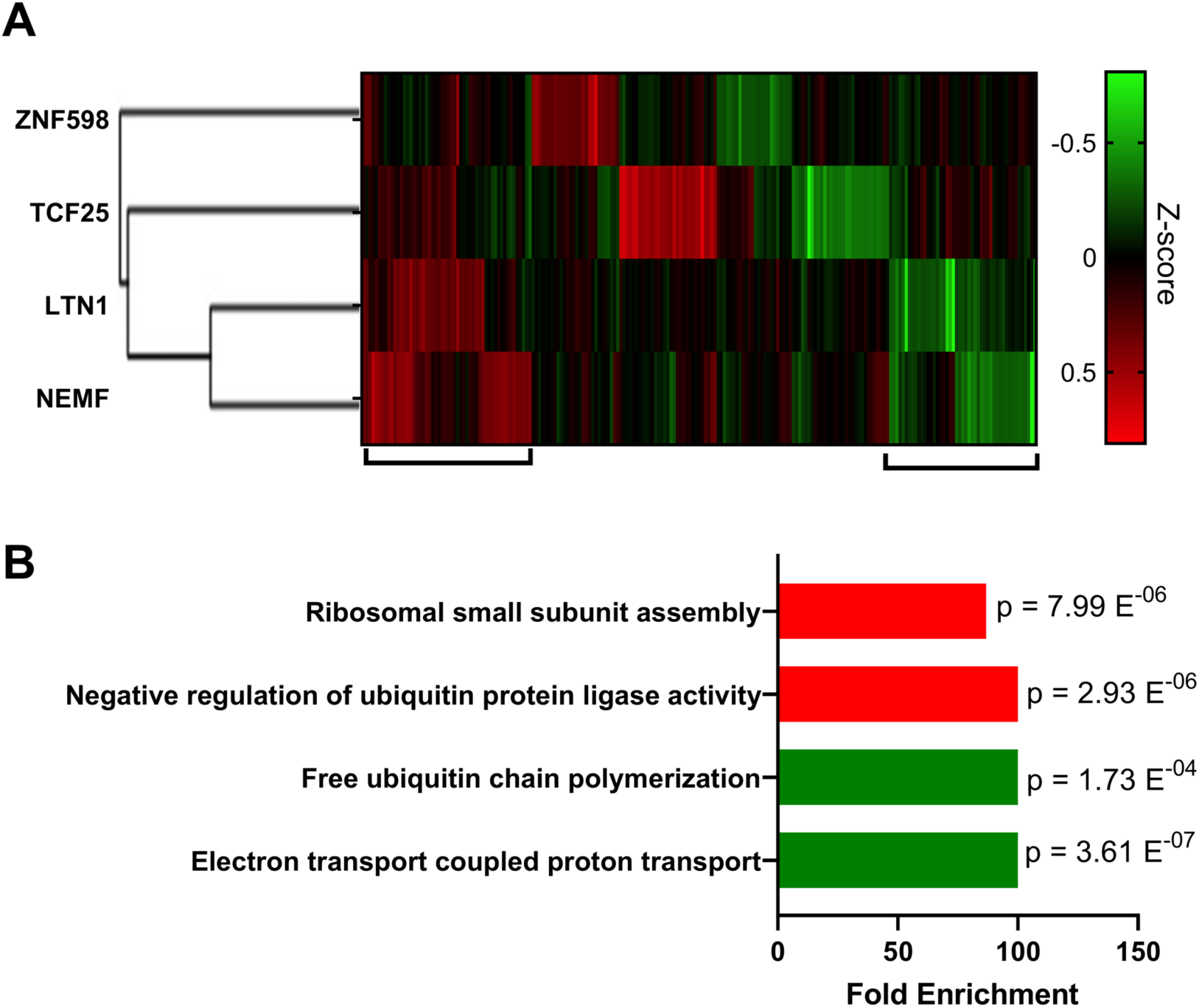
Transcriptional regulation elicited by knockdown of RQC in human cells. (**A**) Heat map of the Z-score distribution of the transcriptional profile when RQC genes were downregulated by CRISPR interference (CRISPRi) in leukemia K562 cell line. Data obtained from Replogle et al., 2022. (**B**) Ontology enrichment of genes that showed the greatest similarity in expression profile when we silenced LTN1 and NEMF, as shown in panel A. The results for up and downregulated gene clusters are shown in **red** and **green**, respectively.

Another interesting feature extracted from the Perturb-seq data is a list of silenced genes presenting a similar transcriptional response. **Figure 6** shows a Venn diagram with the top 10 genes that generated the most similar transcriptional phenotype compared to the knockdown of the RQC components. None of the 4 RQC genes produced a similar transcriptional response (top 10). Moreover, ZNF598 and TCF25 lists showed no overlap whatsoever, but LTN1 and NEMF had 7 hits in common (TMA16, SPATA5L1, HERC1, TEX10, LAS1L, DDX55, and SURF6). Interestingly, none of these 7 genes were previously described in the RQC pathway and have diversified functions with no obvious connections with ribosome quality control or proteostasis **(Supplementary Table 1)**. To check if these genes could participate in the co-translational quality control, we used data from Hickey et al., 2020, which measures the expression of a fluorescence reporter lacking a stop codon (leading to non-stop decay) in a genome-wide CRISPRi screening. As expected, the knockdown of LTN1 and NEMF greatly enhanced GFP fluorescence, while TCF25 and ZNF598 showed smaller but statistically significant results (**Figure 6**). In contrast, none of the 7 genes identified in Perturb-seq analyses (**Figure 6**) affected the GFP fluorescence level, indicating that they are not directly involved in co-translational quality control pathways (**Figure 6**).

**Figure 6.**
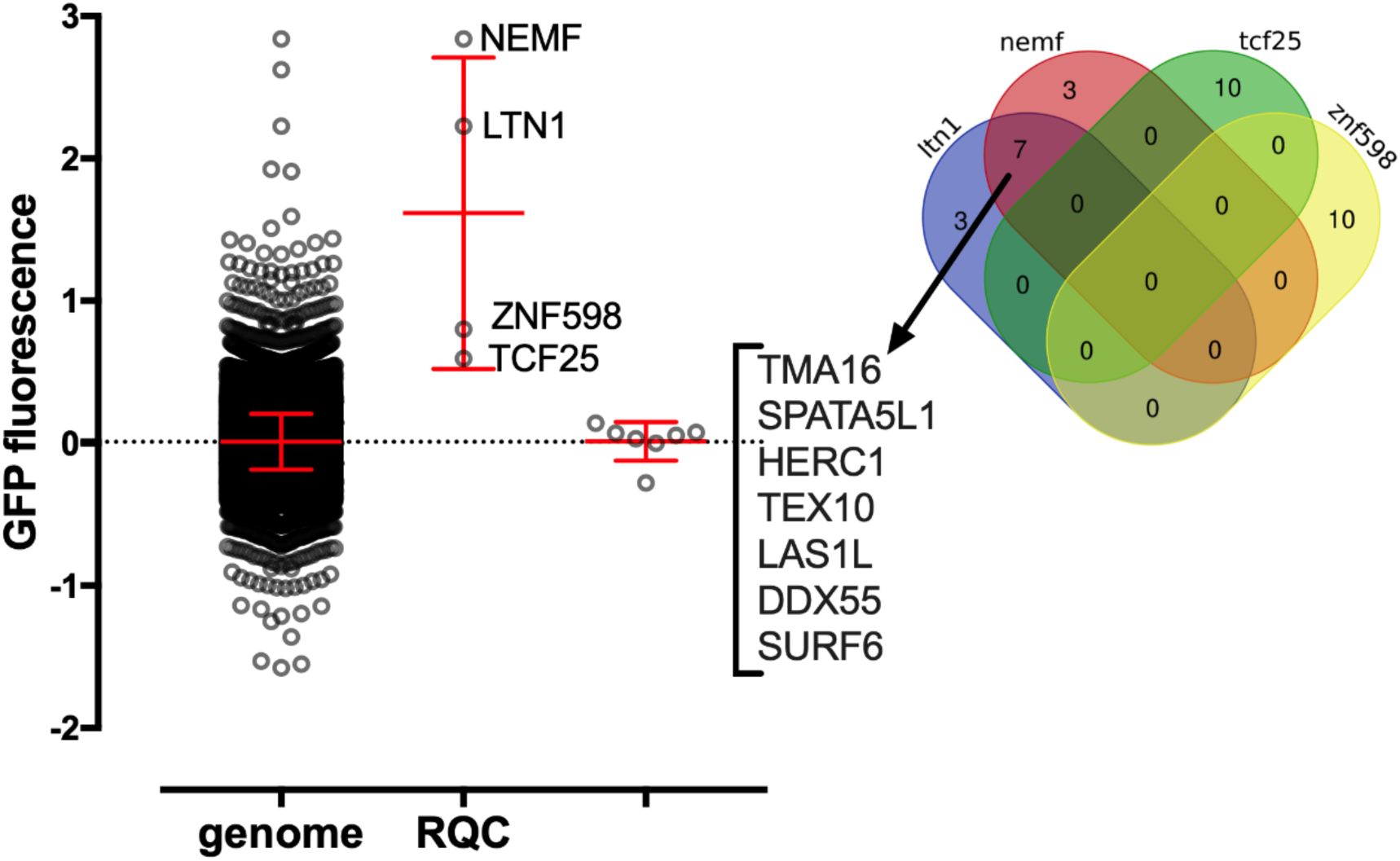
CRISPRi knockdown genes that produced a similar transcriptional phenotype to LTN1 and NEMF are not directly involved in RQC function (Right). Venn diagram shows the top 10 knockdown genes with a similar transcriptional response for each RQC component and ZNF598, of which 7 genes have a similar profile to NEMF and LTN1 knockdown. (**Left**) The graph of GFP fluorescence shows an increase in fluorescence levels when the components of the RQCs complex are deleted, which was not observed for any of the 7 genes with a similar transcriptional profile. Data obtained from.^49^

### The effect of aging on the mRNA levels of the RQC components

Proteostasis failure with the accumulation of insoluble aggregates is a hallmark of several neurodegenerative diseases^56,57^. Failure of the RQC, more specifically of LTN1, can lead to the accumulation of CATylated peptides that promote the formation of protein aggregates due to the CAT tail hydrophobic nature. These aggregates can disrupt neuronal morphogenesis^26,38,58^. Based on this information, we asked if the dysfunction of the RQC contributes to the formation of the insoluble protein aggregates related to old age. To assess if the mRNA levels of the RQC components are affected by aging, we evaluated the transcriptome from several tissues, from donors with different ranges, obtained from the GTEx consortium. We used the transcript per million (TPM) to determine the mRNA abundance across the human tissues at different ages (**Figure 7 A-D**). Compared with the youngest individuals, mRNA levels of the RQC components in aged tissues showed a slight downward trend, which was more evident for NEMF, followed by LTN1 (**Figure 7A-D**). TCF25 also showed a statistically significant decrease (**Figures 7B and 7C**), while no difference was observed for ZNF598 (**Figure 7D**). We did a cluster analysis to analyze tissue-specific age-dependent regulations. NEMF and LTN1 showed a similar pattern of transcriptional regulation. Both genes showed repression in 5 tissues with significant p-values (**Figures 7E and S2)**. TCF25 and ZNf598 showed a more complex and diverse pattern of the regulation **(Figures S3 and S4)**. While there was no statistically significant difference in the brain, we could see a mean reduction trend of the TCF25, LTN1, and ZNF598 mRNA.

**Figure 7.**
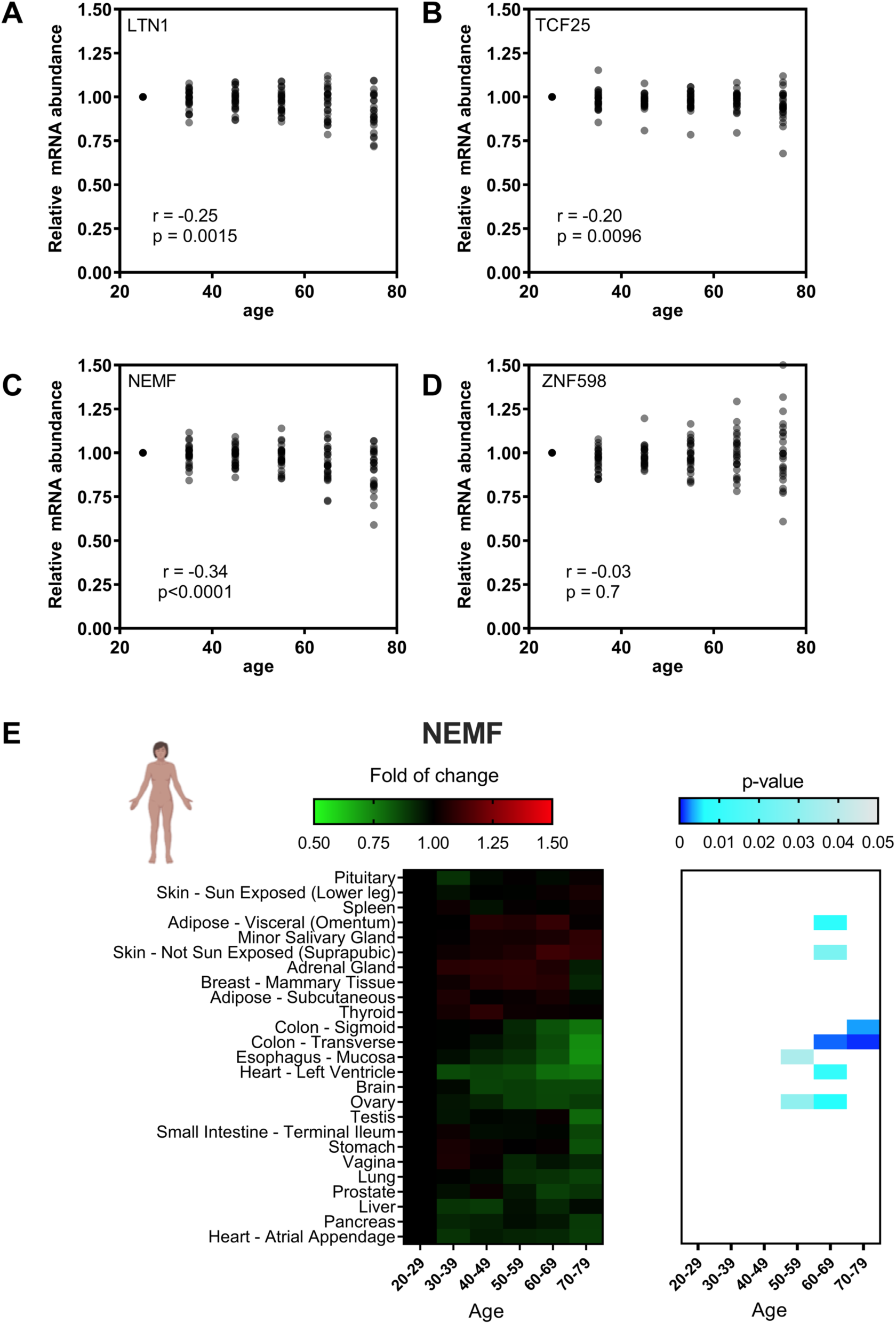
Effects of cellular aging on the transcriptional profile of NEMF, LTN1, TCF25, and ZNF598 in different human tissues. (A-D) Comparison of the relative abundance of mRNA levels in human tissues in age ranges (20-80 years). Data obtained from GTEx consortium. (E) Distribution of the Z-score value of the relative abundance of NEMF mRNA in human organs in different age clusters. Statistical significance was calculated using p-value and presented in a blue gradient for those relevant samples. All data were normalized by the reference age range (20-29).

We sought another model to test the age-dependent regulation observed with GTEx data. We repeated the analyses described in Figure 7 now using data from the *tabula muris* project ^51^, which offers transcriptomes of 20 organs from four male and three female mice ranging from 1 to 27 weeks old. Again, no age-dependent regulation was observed for the RQC components **(Figure S5-8)** probably due the limited number of samples.

## Conclusion

Our results suggest that the RQC complex components are widely expressed in human tissues, suggesting that the RQC is part of the housekeeping processes of the cell **(Figures 1 and 2)**. LTN1 and NEMF share similarities in their expression levels **(Figures 1 and 2)** and their functional and transcriptional phenotypes. However, TCF25, the third essential component of the RQC complex, showed a distinct regulation profile that produced different phenotypes compared with LTN1 and NEMF. This observation can be explained by TCF25 functions outside of the RQC^32–35^.

Our results also showed that TCF25 has a high level of mRNA, low levels of proteins, a high half-life of mRNA, and low half-life of proteins **(Figure 3).** We detect an increase in TCF25 protein levels after blocking the proteasome (**Figure 3D**). These results support a post-translational regulation of TCF25 expression, which has already been experimentally examined in the literature in yeast cells^15^. It has been proposed that the RQC complex regulates Rqc1/TCF25 expression co-translationally through ubiquitination of its polybasic region promoted by RQC activity^25^. However, our group has recently shown that this regulation occurs post-translationally and through the activity of LTN1 alone^15^. TCF25 regulation may impact the RQC complex activity or the transcriptional control exerted by TCF25 in the nucleus. Further studies are necessary to understand how these functions are coordinated and if there are tissue specificities for TCF25 molecular roles.

Further, we observed significant differences in mRNA expression levels of the RQC components when we compared young with aged tissues in humans. A similar effect was observed with *S. cerevisiae* and *C. elegans*^40^. NEMF and LTN1 showed the most consistent age-dependent down-regulation pattern in humans. In *C. elegans*, NEMF and TCF25 homologs showed repression in aged specimens, while in yeast, LTN1, NEMF, and ZNF598 homologs were affected^40^.

We observed a trend of decreased mean expression of the RQC components in the human brain (although not statistically significant), which could be relevant to age-related neuronal impairment **(Figures 7E, S2 and S3)**; ZNF598, contrarily, showed a trend toward increased expression, perhaps as a response to the failure of the RQC **(Figure S4)**. These findings bring insights into the regulation of the RQC complex’s components and its tissue-specific pattern, which could help clarify some phenotypes observed when the activity of the complex is affected by mutations or during aging. A limitation of our study was the comparisons between whole genome data or tissue data, as well as cell data from mouse and human cells. However, we hope that some insights presented herein might be further tested experimentally.

## Acknowledgments

We thank Caio Oliveira and Elis Moll for helpful discussions.

## Author contributions

OALS, RLC, RDR, TD, and FLP were involved in designing and discussing the experiments. OALS, TD, and FLP were involved draft and revision of the paper. OALS was responsible for the western blot. MRA was involved in the statistical analysis of the data.

## Data availability statement

Supplemental data for this article can be accessed on the publisher’s website.

