## Supplemental Figures for "Transcriptional profile of ribosome-associated quality control components and their associated phenotypes in mammalian cells"

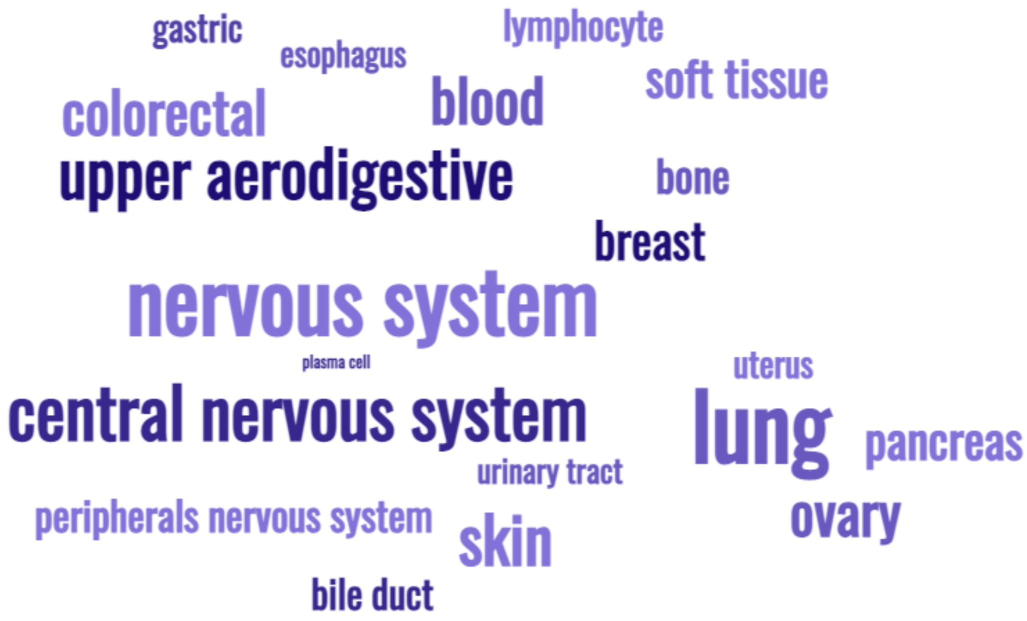


**Supplementary Figure 1.** Word cloud representation of the cellular lineages used to determine the impact of CRISPR-individual gene inactivation for LTN1, NEMF, and TCF25. Data obtained from Tsherniak et al., 2017.

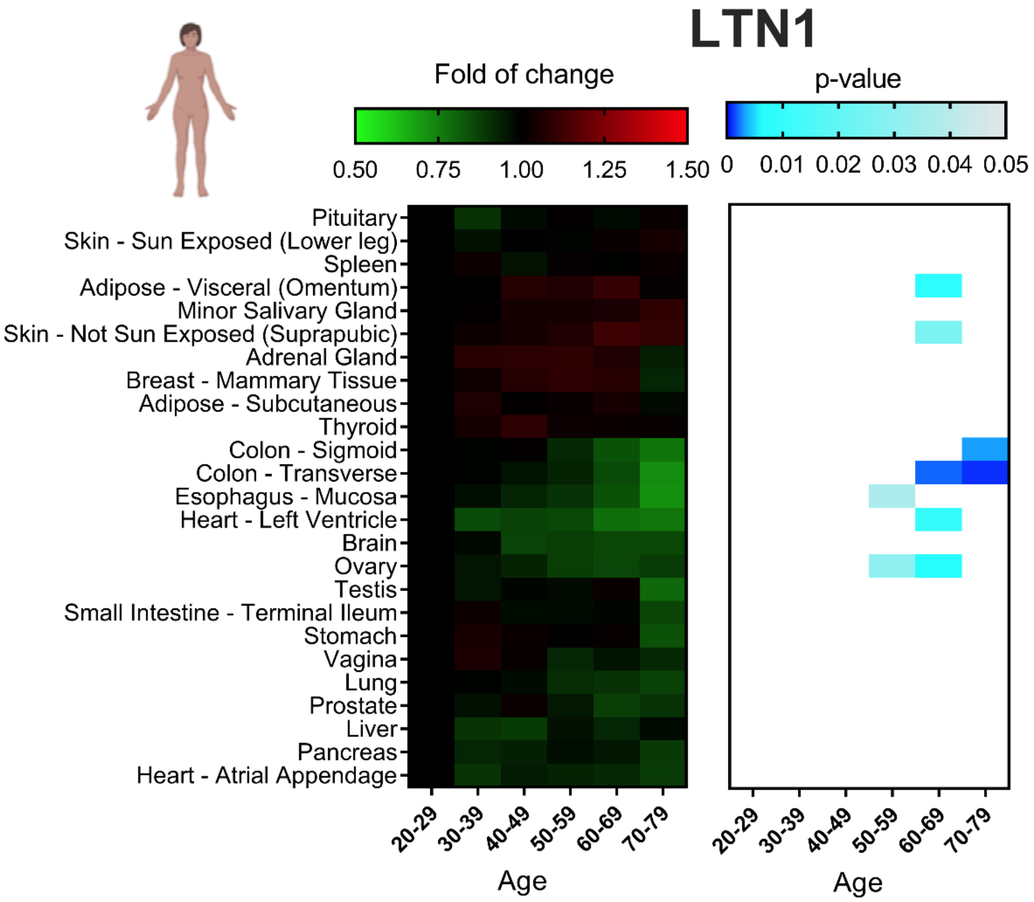


**Supplementary Figure 2.** Distribution of the Z-score value of the relative abundance of LTN1 mRNA in main human organs in different age clusters. Statistical significance was calculated using p-value and presented in a blue gradient for those samples that showed relevance. All data were normalized by the reference age value (20-29).

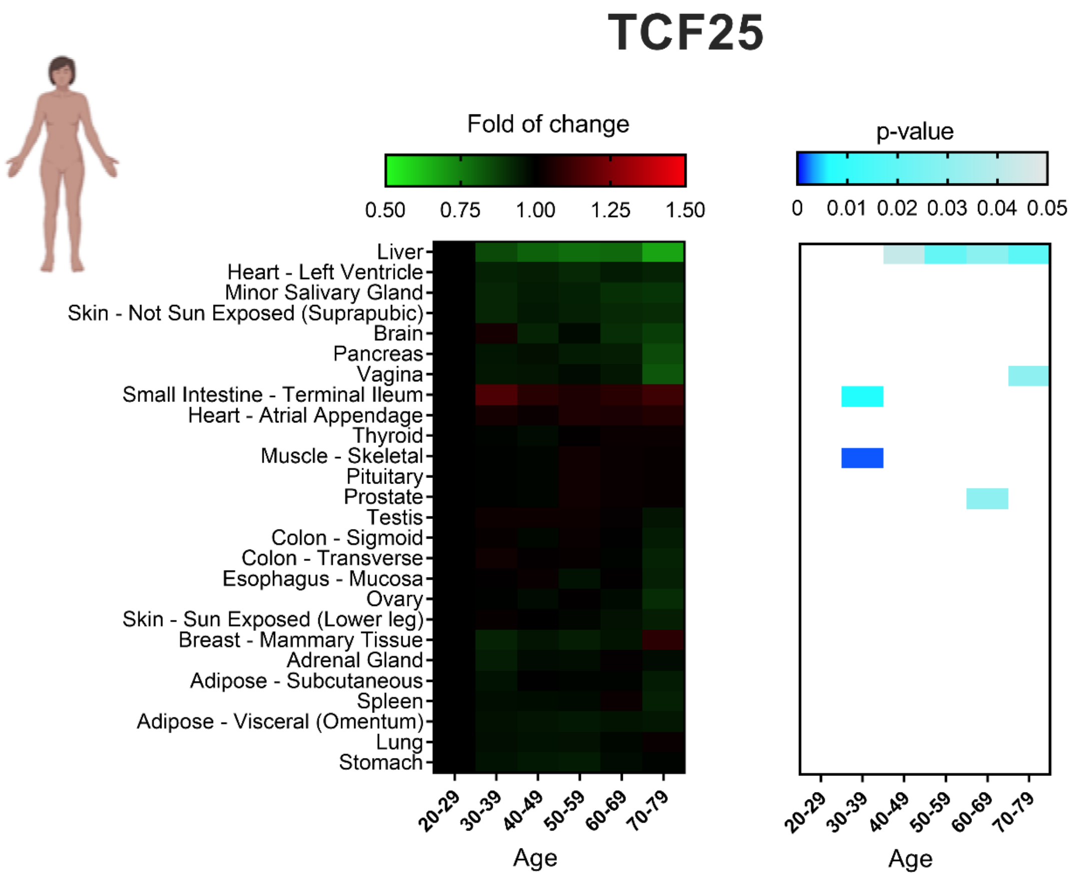


**Supplementary Figure 3.** Distribution of the Z-score value of the relative abundance of TCF25 mRNA in main human organs in different age clusters. Statistical significance was calculated using p-value and presented in a blue gradient for those samples that showed relevance. All data were normalized by the reference age value (20-29).

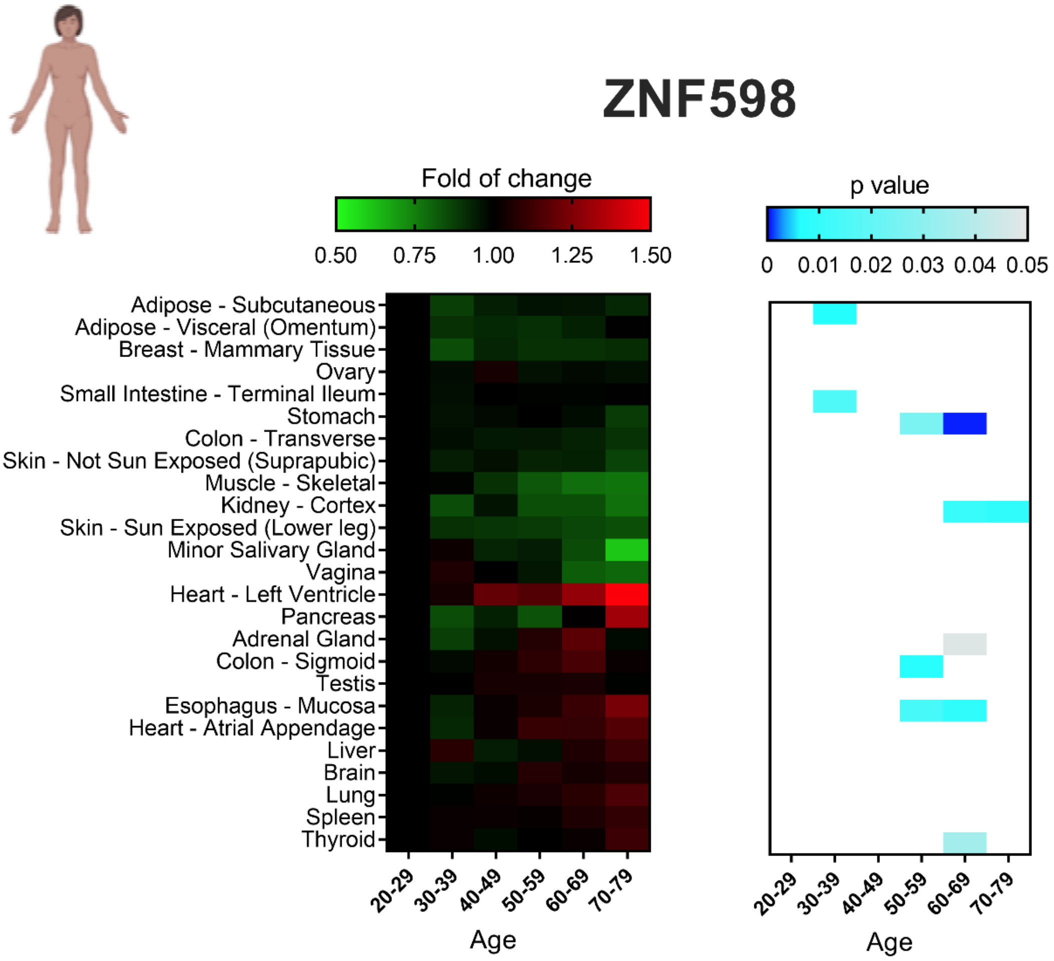


**Supplementary Figure 4.** Distribution of the Z-score value of the relative abundance of ZNF598 mRNA in main human organs in different age clusters. Statistical significance was calculated using p-value and presented in a blue gradient for those samples that showed relevance.

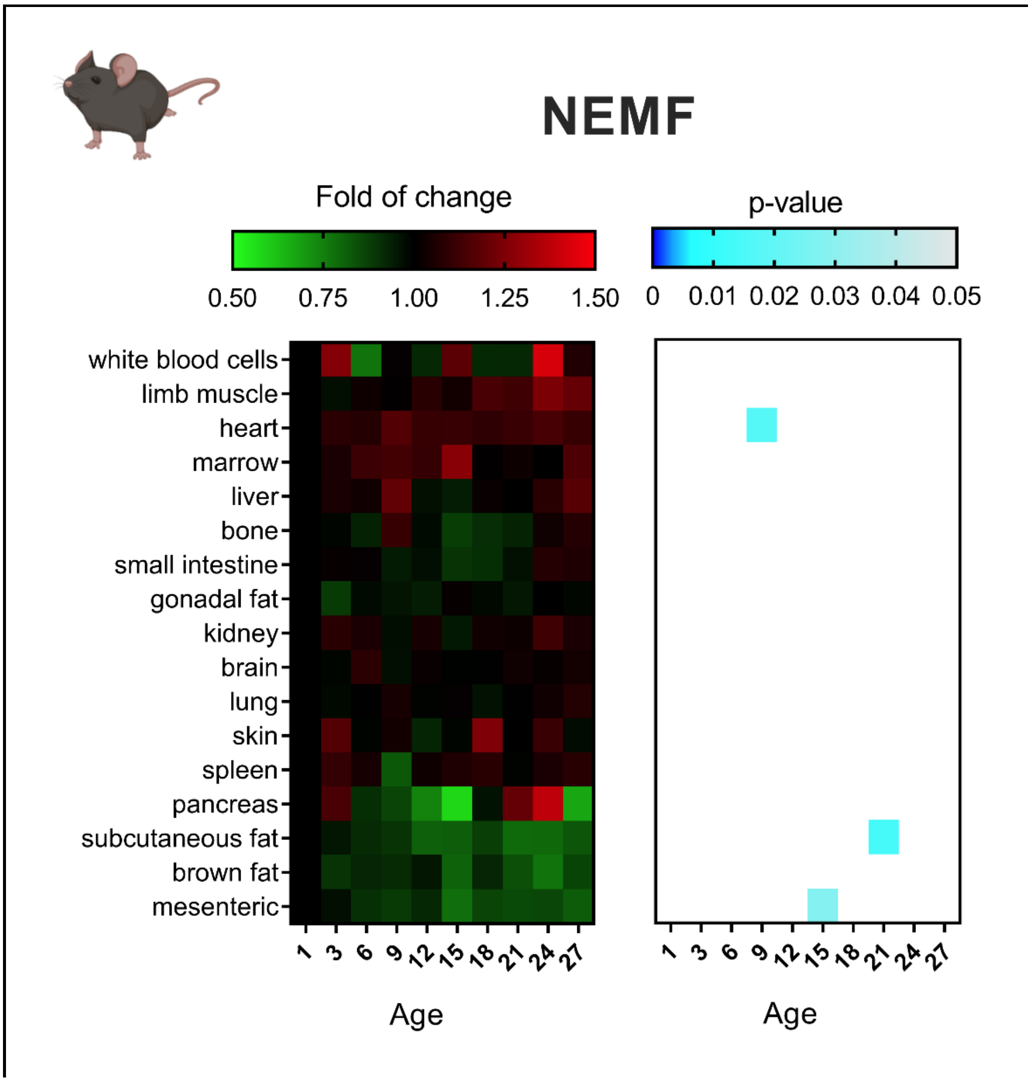


**Supplementary Figure 5.** Distribution of the Z-score value of the relative abundance of NEMF mRNA in main mouse organs in different age clusters. Statistical significance was calculated using p-value and presented in a blue gradient for those samples that showed relevance. All data were normalized by the reference age value (1 week).

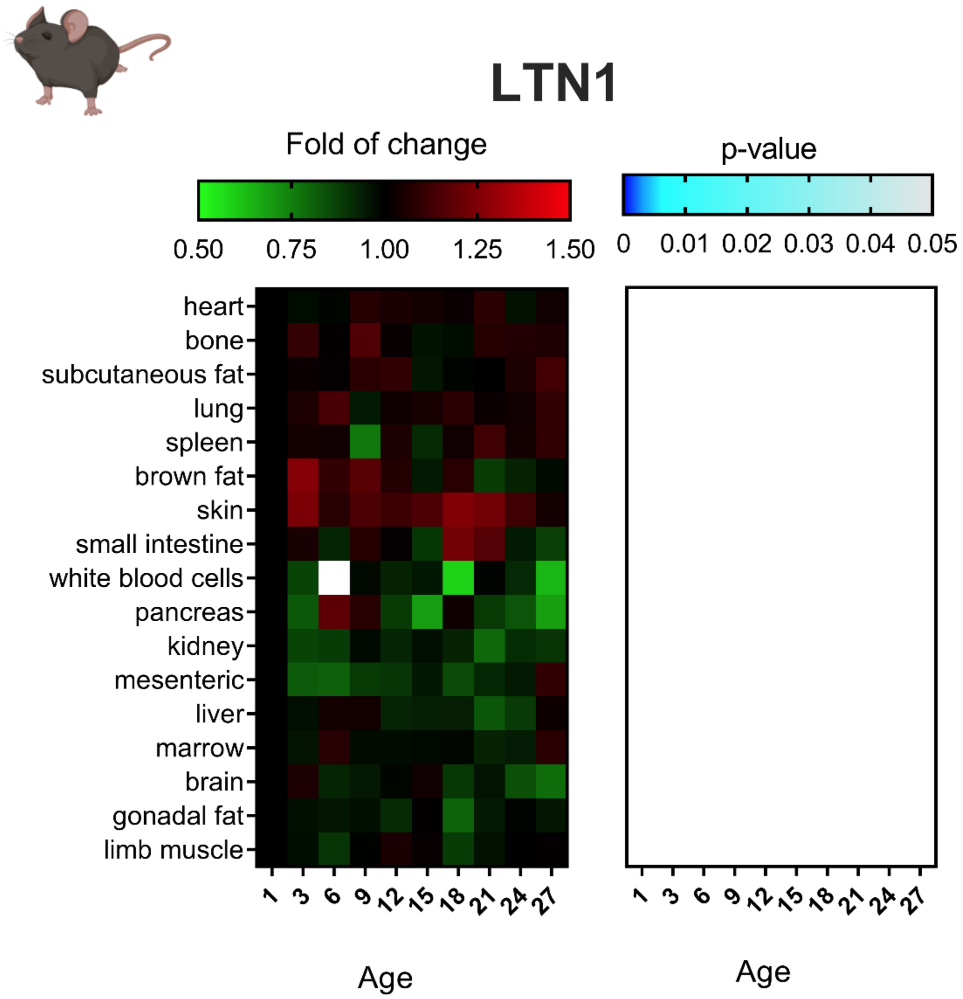


**Supplementary Figure 6.** Distribution of the Z-score value of the relative abundance of LTN1 mRNA in main mouse organs in different age clusters. Statistical significance was calculated using p-value and presented in a blue gradient for those samples that showed relevance. All data were normalized by the reference age value (1 week).

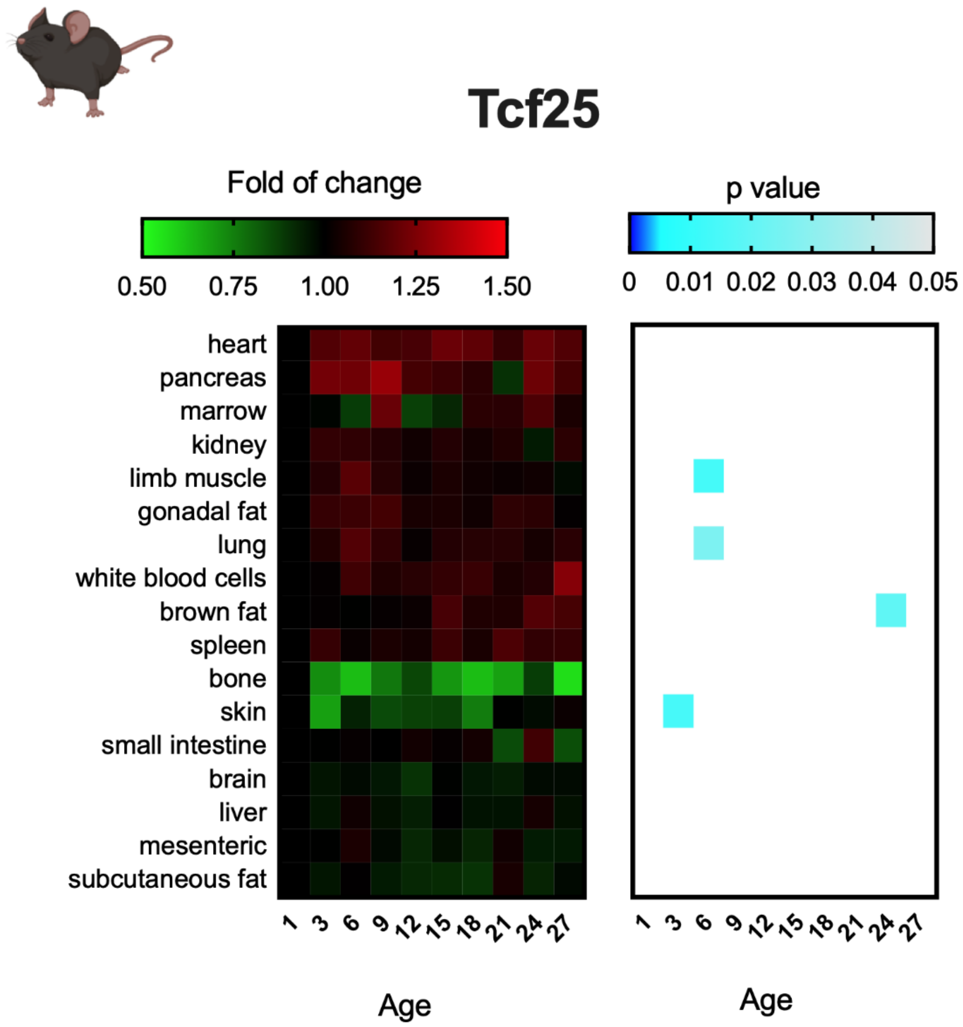


**Supplementary Figure 7.** Distribution of the Z-score value of the relative abundance of TCF25 mRNA in main mouse organs in different age clusters. Statistical significance was calculated using p-value and presented in a blue gradient for those samples that showed relevance. All data were normalized by the reference age value (1 week).

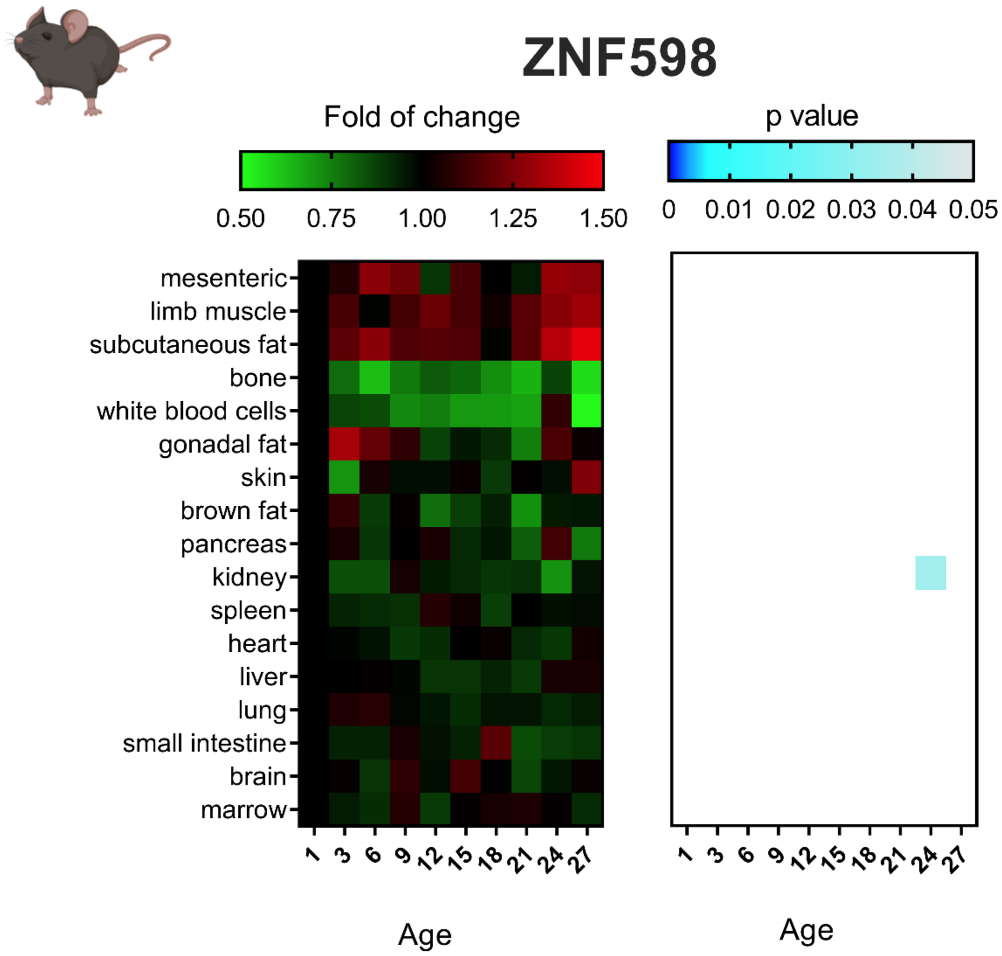


**Supplementary Figure 8.** Distribution of the Z-score value of the relative abundance of ZNF598 mRNA in main mouse organs in different age clusters. Statistical significance was calculated using p-value and presented in a blue gradient for those samples that showed relevance. All data were normalized by the reference age value (1 week).

**Supplementary Table 1**: Gene function of the 7 genes with similar transcriptional profile with NEMF and LTN1 obtained from Ensembl.

| Gene name | Ensembl ID | GO: Biological process |
|---|---|---|
| TMA16 | ENSG00000198498 | Ribosome biogenesis |
| SAPTA5L1 | ENSG00000171763 | Ribosome biogenesis |
| HERC1 | ENSG00000103657 | Autophagy<br>Brain development Protein ubiquitination Cerebellar Purkinje cell differentiation<br>Corpus callosum development<br>Neuron projection development<br>Regulation of catalytic activity<br>Neuromuscular process controlling balance |
| TEX10 | ENSG00000136891 | Not available |
| LAS1L | ENSG00000001497 | rRNA processing and maturation<br>Nucleic acid phosphodiester bond hydrolysis |
| DDX55 | ENSG00000111364 | Not available |
| SURF6 | ENSG00000148296 | Ribosome biogenesis |
